# Advanced Maternal Age Impairs Decidual Angiogenesis and Spiral Artery Remodelling

**DOI:** 10.64898/2026.10.21.760773

**Authors:** Wendy K. Duan, Natalie Nakatsui, Alyssa Spyracopoulos, Elmy Castillo, Juan Garcia Rivas, Christian Bellissimo, Meghan Riddell

## Abstract

Pregnancy presents an extreme physiological challenge that relies on rapid angiogenic vascular expansion and remodelling to establish the maternal-fetal vascular unit at the uteroplacental interface. These adaptations to the pregnancy-specific decidual vasculature are necessary for pregnancy progression. Advanced age pregnancies (AAP; pregnant person >35 years old) are associated with increased risk of poor pregnancy outcomes. Despite increasing evidence of the centrality of vascular aging in mediating organotypic dysfunction, whether vascular aging in the decidua occurs in the context of AAP is unknown. Using a murine model of AAP, we find that alongside embryonic developmental delay and deficient placental vascularization, AAP dams had impaired decidual angiogenesis and spiral artery remodeling with imbalances in both systemic and local uterine angiogenic factor expression in the VEGF and angiopoietin signalling families. These findings reveal that pregnancy may expose vascular aging mechanisms that contribute to declining reproductive success and increased cardiovascular risk later in life.

## Introduction

Pregnancy has been referred to as the ultimate cardiovascular stress test since complicated pregnancies have been associated with an increase in maternal cardiovascular disease later in life.^1–4^ Pregnancy poses a unique cardiovascular challenge as it requires rapid and dynamic adaptations in uterine and systemic vasculature. A primary site of vascular adaptation occurs when the endometrial lining transforms into the pregnancy-permissive decidua. During this transition angiogenic expansion and remodelling of decidual vessels establish the maternal-fetal vascular unit. Beginning post-implantation in the murine decidua, VEGF- and angiopoietin-driven^5–10^ angiogenesis and vascular enlargement results in a global increase in vascular diameter and density by mid-gestation.^7,8^ From gestational day (GD) 10.5 onwards, decidual vasculature continues to increase in diameter and undergo morphological changes, coinciding with the accelerated growth of the placenta.^11^ Alongside these global changes, starting from mid-gestation spiral arteries are transformed into high-capacitance, low-resistance vessels, necessary for the establishment of healthy uteroplacental circulation. Driven by endothelial-intrinsic and -extrinsic mechanisms, spiral arterial vessel diameters expand and the muscular sheathe and extracellular matrix (ECM) surrounding the endothelium are lost.^12–15^ Critically, defects in vascular remodelling or aberrant uterine vascular structures have been associated with adverse pregnancy outcomes such as stillbirth, miscarriage, preeclampsia, and fetal growth restriction.^10,16–20^

Advanced age pregnancy (AAP), otherwise defined as advanced maternal age, is when a pregnant person is over the age of 35. AAPs are at a higher risk of adverse pregnancy outcomes including preeclampsia, gestational hypertension, fetal growth restriction, and stillbirths.^21–27^ Internationally, AAPs constitute a significant and growing patient population.^28,29^ Along with decreasing oocyte quality, the aging uterus is a major contributor to age-associated risk during pregnancy. Woods *et al.* (2017) showed that many AAP-associated outcomes were mitigated upon AAP embryo transfer to young uteri, demonstrating the importance of the uterine environment in determining pregnancy outcomes with age.^30^

Vascular attrition and microvascular rarefication are hallmarks of tissue-specific aging, resulting in impairments in the ability of endothelial cell (EC) to mount angiogenic responses.^31–37^ Across multiple organs, age-associated angiogenic impairments are due to both EC-extrinsic microenvironmental factors and EC-intrinsic factors. In aging mice, angiogenic VEGF signalling insufficiency arises as a result of both increases in systemic VEGF-antagonistic sFLT1 and decreases in tissue-localized VEGF.^38,39^ Furthermore, it is well established in other organs that EC exhibit a high degree of organotypic heterogeneity^40,41^ and age in an organotypic manner.^31,32,34^ Despite this increased appreciation of angiodiversity and its intersection with age, little is known concerning the aging decidual vasculature. While AAP is known to alter maternal cardiac function, peripheral vascular resistance, and uterine artery function in humans and rodents,^42^ no previous research has characterized its impact on decidual vascularization and spiral artery remodelling. Thus, whether the aging uterine vasculature impacts the ability of older individuals to mount the necessary angiogenic responses during pregnancy remains unknown. Given the importance of decidual angiogenesis during pregnancy, vascular aging may be a key understudied mechanism mediating declining uterine function and increased AAP-associated adverse pregnancy outcome risk.

Here, using a murine model of AAP, we characterize the decidual vasculature in young vs. AAP dams and find that, alongside severe impairments in fetal and placental development, there is impaired vascular development and spiral artery remodelling in AAP deciduae and placentae. Interestingly, there are imbalances in maternal circulating and local decidual angiogenic factor expression, suggesting the impaired development of decidual vasculature with AAP may be driven by tissue-specific and systemic impairment of angiogenic signalling.

## Results

### AAP leads to embryonic developmental delay and diminished placental vascularization

To establish whether defects in the angiogenic expansion and remodelling of the decidua are a feature of AAP, we developed a murine AAP model where young (11-16wks) and AAP (39-69wks) C57BL/6J females were mated with young (11-16wks) C57BL/6J males. Increasing age is associated with significant changes in body composition and weight gain in humans.^43–45^ To mitigate excessive weight gain within our aged cohorts, females were pair fed starting at 20wk. Prior to mating, aged females were heavier (27.3±2.8g) than young dams at 11wks (22.15±1.6g) (Supplementary Figure 1A). This increase in weight was contributed by an increase in both fat and lean mass, with greater gains in fat mass (Supplementary Figure 1B-D). Notably, aged mice were not obese, and the increased weight and adiposity was considered within the range of expected age-associated weight gain.^43^ Thus, our aging model represents a non-obesogenic maternal aging phenotype.

Dams were mated then sacrificed at GD10.5 (mid-gestation) when maternal circulation to the placenta has begun^46–48^ and spiral artery remodelling is occurring.^11^ To assess whether hormone production was altered in AAP dams, maternal plasma was assessed for circulating steroid hormones (Supplementary Figure 1E-H), where there were critically no differences in circulating estradiol or progesterone. When examining intact uterine horns, the number of implantation sites decreased with increasing maternal age (Figure 1A, B). Uterine horns from AAP dams aged 52-69wk (12+mo) had increased frequency of small, dark, hemorrhagic implantations indicative of active resorptions (Figure 1A). Consistent with this, no embryos nor placentae were recoverable from these hemorrhagic implantation sites. These active resorptions were not prevalent in young dams nor AAP dams aged 39-52wk (9-12mo). Although the uterine horns of AAP dams aged 9-12mo appeared grossly similar to young dams, dramatic differences in embryonic phenotypes could be observed. Embryonic morphology was assessed from 6 young and 6 AAP dams using Theiler staging.^7^ Compared to control embryos, embryos from AAP dams had a wider range of developmental variability and were more likely to be developmentally delayed even within the same litter (Figure 1C, D). While 89.4% of control embryos were developmentally appropriate at GD10.5, only 29.0% of embryos from AAP dams were appropriate, while the rest were delayed by 1-1.5 or 2+ gestational days (Figure 1D). When comparing developmentally normal embryos from young and AAP litters, there was no difference in crown rump length at GD10.5 (Figure 1E), hence there is no discernible growth restriction between embryos of a similar developmental stage at this gestational stage.

**Figure 1:**
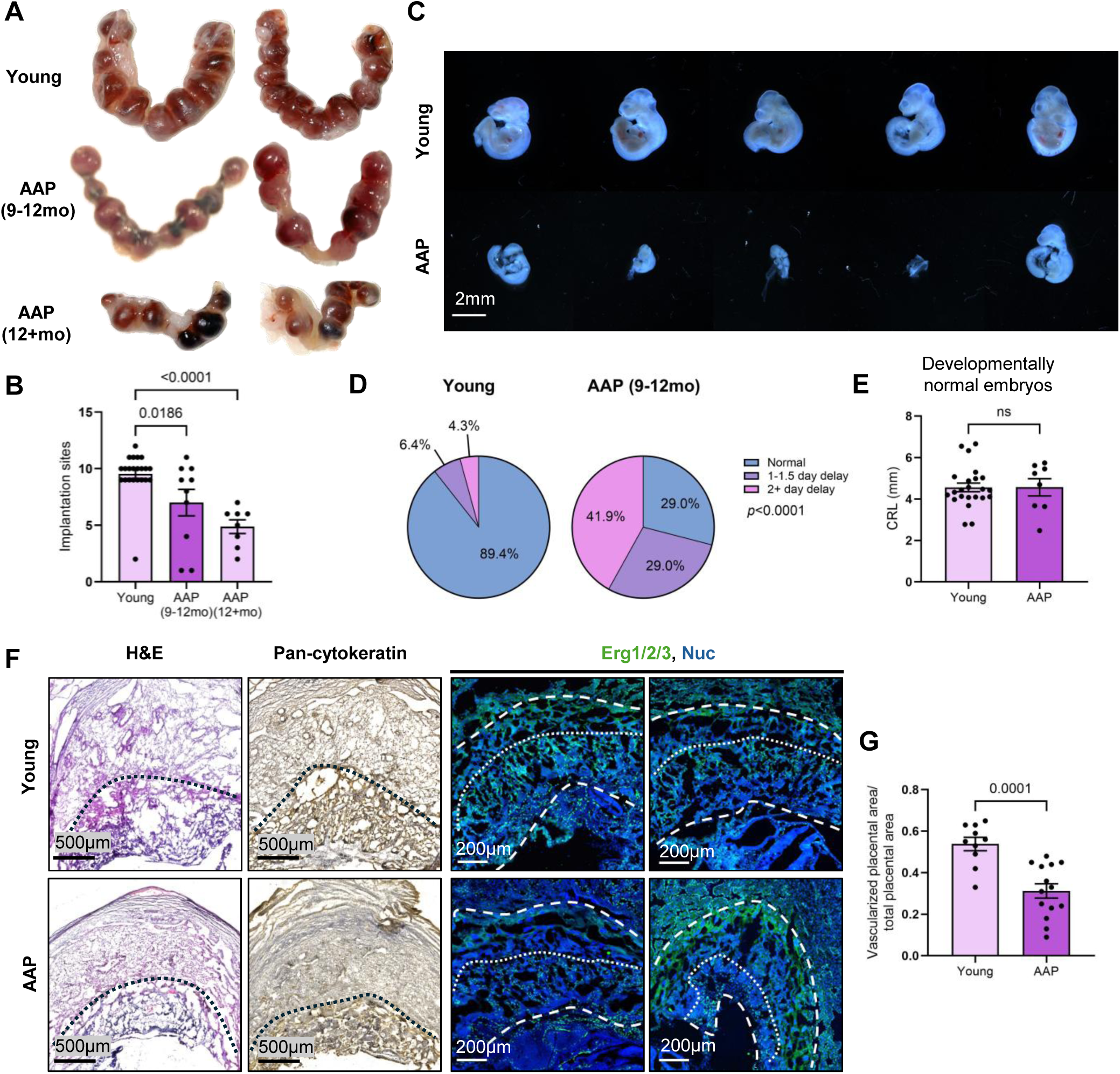
Embryos from AAP dams at GD10.5 are more likely to be developmentally delayed and have altered placental composition. A) Representative images of whole uterine horns at GD10.5; B) Summary data of number of implantation sites at GD10.5; C) Representative stereomicroscopy images of embryos at GD10.5; D) Summary data of embryo development stages at GD10.5 in young vs AAP dams; E) Summary data of crown rump length (CRL) of developmentally normal embryos in young vs AAP dams; F) Representative images of implantation sites stained for H&E, anti-pan-cytokeratin, and anti-ERG1/2/3 and Hoechst at GD10.5; big dashed lines = placental area, small dashed lines = upper boundary of vascularized placenta; G) Summary data of the vascularized placental area as a fraction of total placental area; B) mean ± SEM, one-way ANOVA with multiple comparisons; D) chi-squared test; E, G) mean ± SEM, student’s t-test; 47 embryos from 6 young dams and 31 embryos from 6 AAP dams were analysed.

To assess placental morphology, intact implantation sites from young and AAP dams were visualized using hematoxylin and eosin (H&E) staining and immunostaining with anti-pan-cytokeratin (trophoblastic marker) and anti-Erg1/2/3 (pan-EC nuclear marker) (Figure 1F). Placentae from AAP dams had diminished vascularization, suggesting impaired labyrinth zone development, critical for nutrient and gas exchange between maternal and fetal circulations (Figure 1F, G). Consistent with Woods *et al.* (2017)^30^ and our structural analyses, RT-PCR analyses (Supplementary Figure 2) revealed that AAP placentae exhibited upregulated expression of stem-like trophoblast (*Eomes, Esrrb, Elf5*) and trophoblast giant cell (TGC; *Prl2c2*) marker genes alongside downregulation of intermediate trophoblast (*Tpbpa*) and labyrinthine (*Gcm1, Synb, Ctsq*) marker genes, indicating expansion of stem-like trophoblasts and TGC at the expense of the labyrinthine zone. Overall, our AAP model results in increased embryonic loss post-implantation, an increased incidence of embryonic developmental delay, and diminished placental vascularization.

### Spiral artery remodelling is impaired in AAP

Spiral artery remodelling is marked by arterial lumen dilation with reduced ECM and smooth muscle cell (SMC) coverage,^12,13,49^ ultimately resulting in a thinner vessel wall with reduced musculature and elasticity and a larger lumen through which blood can transit. To compare spiral artery remodelling in young and AAP dams, histological analyses were performed on midsagittal GD10.5 uterine sections (Figure 2A-F). Using Masson’s Trichrome stain, we stereologically analyzed 6-11 implantation sites from 3 young and 3 AAP dams for average lumen/vessel diameter as a modification of Kieckbutsch *et al.*’s method of assessing spiral artery remodelling progression.^49^ AAP implantation sites had impaired spiral artery remodelling, characterized by a 13.1% decrease in average lumen/vessel ratio (Figure 2B). To quantify SMC presence, 7-10 implantation sites from 3 young and 4 AAP dams were co-stained with anti-CD31 and anti-α-smooth-muscle-actin (αSMA) and αSMA coverage was calculated as a function of anti-αSMA sum signal normalized by vessel volume. Spiral arteries from young implantations tended to have thin, sparse SMC, while AAP spiral arteries were smaller and surrounded by a thicker layer of SMC that more fully encapsulated the vessel (Figure 2C). Average αSMA coverage increased 68.4% in AAP implantations vs young, further supporting our conclusions that there is a significant impairment in spiral artery remodelling in AAP dams.

**Figure 2:**
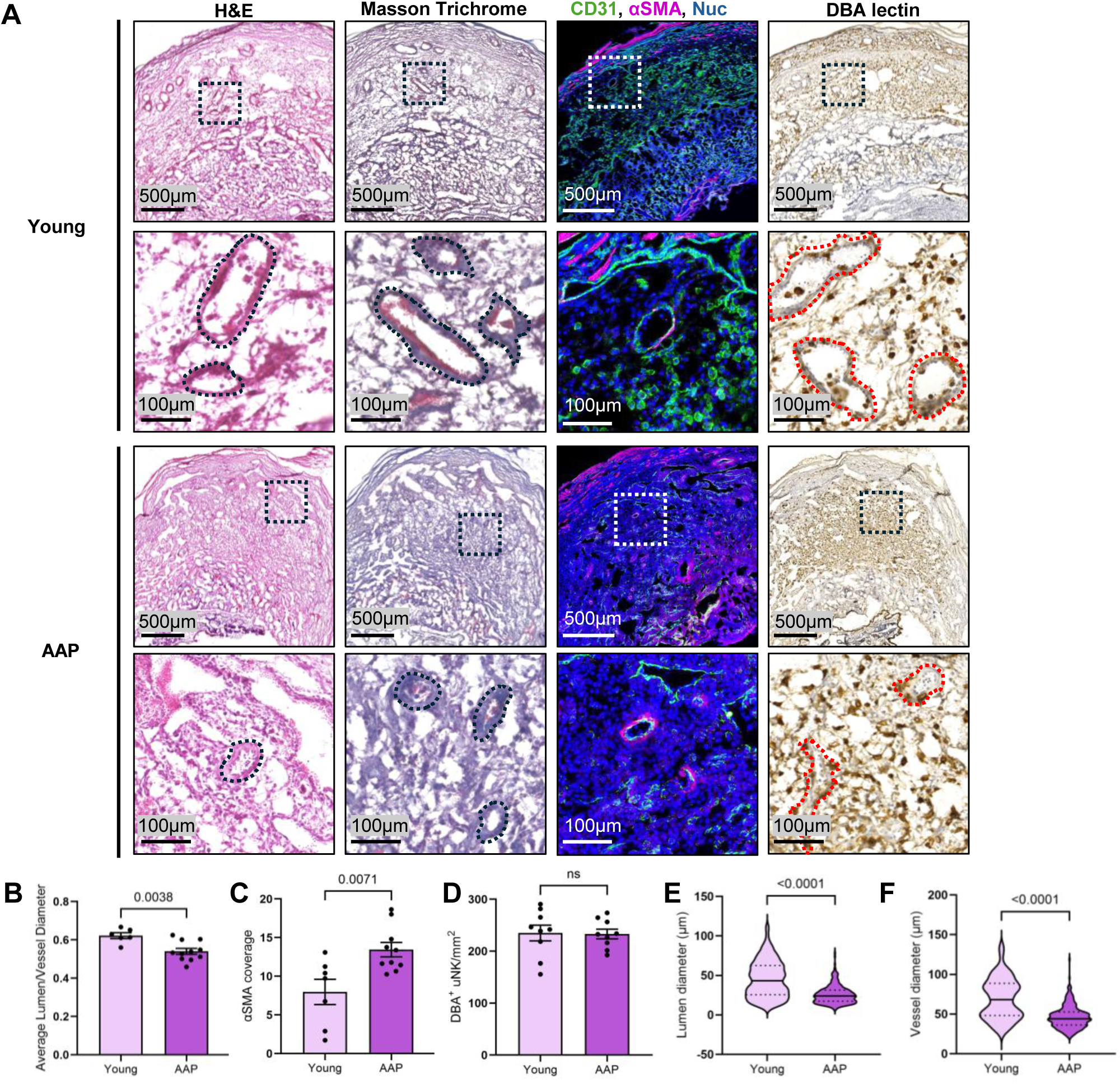
Spiral artery remodelling is impaired in AAP dams. A) Representative images of young vs AAP implantation sites at GD10.5 stained for H&E, Masson’s Trichrome, anti-αSMA and anti-CD31, and DBA lectin; Summary data of B) average lumen-vessel ratio, C) average αSMA coverage, D) DBA^+^ uNK per mm^2^; Violin plots of E) lumen diameter (μm) and F) vessel diameter (μm); B-D) mean ± SEM, student’s t-test; E-F) unpaired Mann-Whitney test; solid line indicates the median; dashed lines indicate quartiles 1 and 3; 6-11 implantations from 3 young and 4 AAP dams were quantified.

In mice, uterine natural killer cells (uNKs) secrete cytokines, angiogenic factors, and dedifferentiation factors that contribute to the disruption of SMCs and breakdown of ECM.^12,13,50–52^ As such, transgenic mice with depletions in DBA^+^ uNKs have impaired spiral artery remodelling.^52,53^ As expected, DBA^+^ uNKs were abundant in the decidual stroma and near vessel walls (Figure 2A). However, there was no change in the abundance of DBA^+^ uNK in the decidua between young and AAP dams (Figure 2D). The role the EC play in early stages of spiral artery remodelling remains poorly understood. Critically, dilation of the spiral arteries can precede SMC removal.^14^ To assess arterial dilation, we compared lumen and vessel diameters between young vs AAP implantations. AAP implantations had a 19.8±11.2μm decrease in lumen diameter and a 23.4±14.5μm decrease in vessel diameter compared to young implantations (Supplementary Figure 3A-B). Both arterial lumen and arterial vessel diameters in young implantations had a bimodal distribution (Figure 2E, F). Comparatively, arteries in AAP implantations for both lumen and vessel diameter lost this bimodal distribution and were largely constrained to the lower half of the respective young distributions. Thus, spiral arteries in AAP implantations failed to dilate, resulting in significantly decreased lumen and vessel diameters and suggesting an endothelial insufficiency.

### Angiogenic vascular expansion is impaired in AAP deciduae

As previously mentioned, the decidual vasculature expands and establishes contiguous vascular connections with the developing placental circulation via angiogenesis, increasing in calibre as pregnancy progresses.^7,8^ To visualize the decidual vasculature, we stained midsagittal uterine sections with anti-CD31, anti-Erg1/2/3, and Hoechst (Figure 3A, B). Vesselucida was used to trace the decidual vasculature in 7-12 implantation sites from 4 young dams and 7 AAP dams from confocal tile scanned images (Figure 3C). These tracing maps allowed us to characterize the structure of this vascular bed, excluding the myometrial and placental vessels. Consistent with arterial analyses (Figure 2E-F), AAP implantations had a global shift towards smaller calibre vessels compared to young (Figure 3B-D). While 41.0% of vessels in young deciduae were ≤10μm, 55.0% of vessels in AAP deciduae were ≤10μm (Figure 3D). This increase in small calibre vasculature was at the expense of larger vessels, particularly those ≥30μm in diameter (Figure 3D). Hence vascular enlargement is severely impaired in AAP decidua. AAP implantations additionally had a 56.6% decrease in decidual vascular density compared to young, measured as a function of vascular volume divided by total decidua volume (Figure 3E). To assess EC abundance, we quantified the percentage of Erg1/2/3^+^ nuclei to total nuclei from randomly selected decidual regions of interest. Compared to young implantations, AAP implantations had a 7.6±5.4% decrease in the proportion of EC per area (Figure 3B, F). Although there are proportionally fewer EC per area in AAP implantation sites, there was no change in the percentage of proliferative Ki67+ EC to total EC at GD10.5 (Figure 3G, H). Therefore, these decreases in vascular diameter, EC abundance, and vascular density demonstrate that by mid-gestation AAP decidua have impaired angiogenic vascular remodelling but no alteration in EC proliferation.

**Figure 3:**
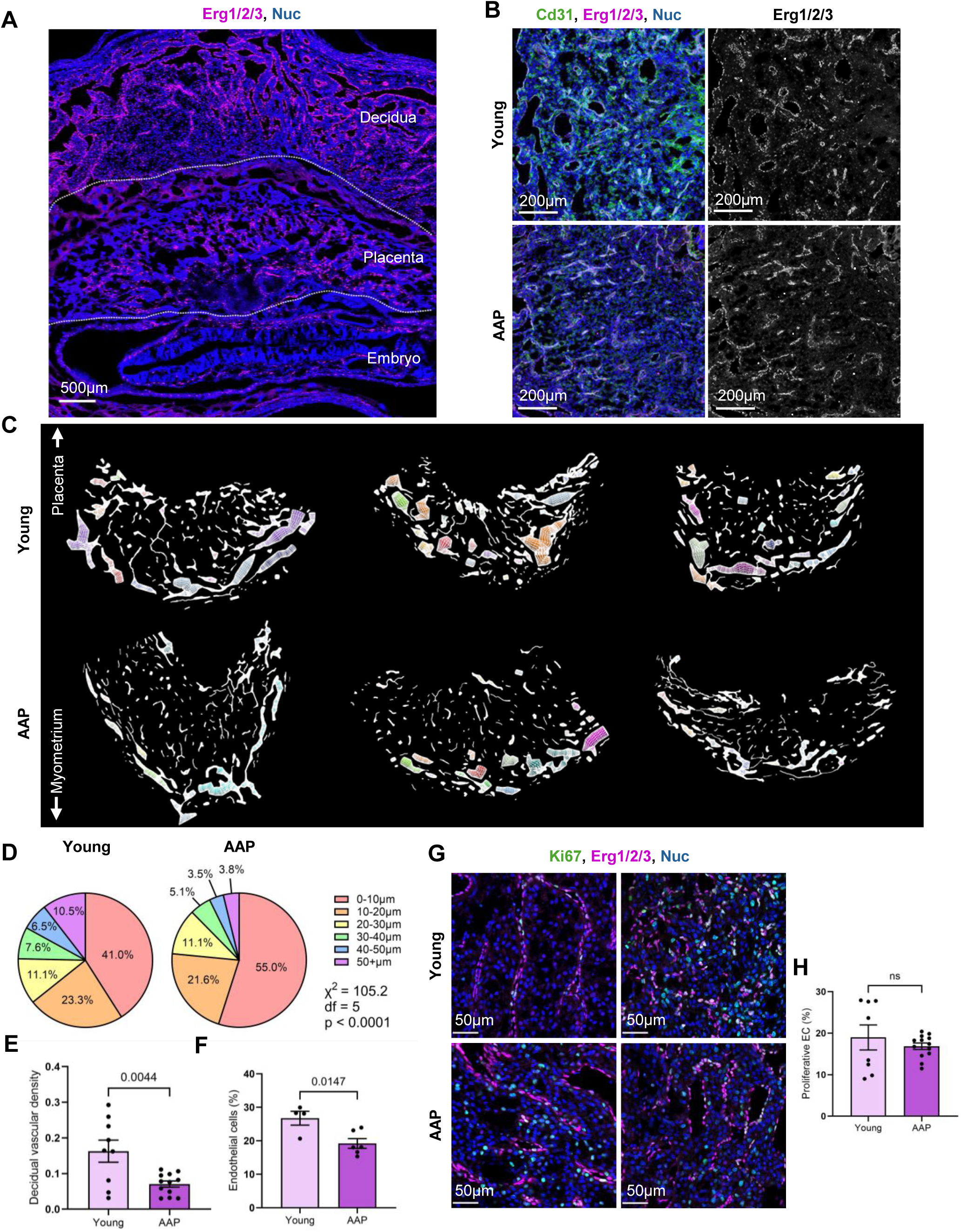
Decidual vascular remodelling at GD10.5 is impaired in AAP dams. A) Representative image of a murine implantation site stained for anti-Erg1/2/3 and Hoechst, outlining the decidua, placenta, and embryo; dashed lines = division lines between the decidua, placenta, and embryo B) Representative confocal microscopy images of murine decidua stained for anti-CD31, anti-ERG1/2/3, and Hoechst; C) Representative vessel tracing maps of decidual vasculature; Summary data for D) distribution of vessel diameters in young vs AAP dams; E) proportion of endothelial cells (%), and F) vascular density; G) Representative image of young vs AAP decidua stained for anti-ERG1/2/3, anti-KI67, and Hoechst; H) Summary data of the percentage of KI67+ endothelial cells (%) in young vs AAP dams; D) chi-squared test; E-F, H) mean ± SEM, student’s t-test; 4-12 implantations from 4 young and 7 AAP dams were quantified.

Since both placental and decidual vascularization was diminished in AAP (Figure 1G, 3E), we assessed whether placental vascularization and decidual vascularization were correlated using paired placental and decidual data. Using a linear mixed-effects model, we tested whether decidual vascular density and maternal age were predictors for placental vascularized fraction, nesting implantations from the same dam (Supplementary Figure 4A). Decidual vascular density was not a statistically significant predictor of placental vascularized fraction in this model. When testing the inverse relationship, or whether placental vascularized fraction and maternal age are predictors of decidual vascular density, placental vascularized fraction was likewise a non-significant predictor of decidual vascular density (Supplementary Figure 4B). Consistent with previous findings in this study, advancing maternal age was a significant predictor and inversely related with both placental vascularity and decidual vascularity (Supplementary Figure 4A-B). These findings show that although these two pregnancy-specific tissues develop in conjunction, vascularity of one is weakly dependent or independent of the other by mid-gestation.

### Imbalances in angiogenic VEGF expression are observed in AAP dams

It is known that angiogenic VEGF signalling is necessary for decidual vascularization^8^ and that VEGF crosstalk between the decidua and the placenta affects both pregnancy progression and maternal cardiovascular function.^54,55^ To elucidate the mechanisms driving impaired angiogenic remodelling, we examined angiogenic factor expression. To assess local production of angiogenic VEGF factors in AAP, we performed RT-PCR on dissected young and AAP deciduae and placentae at GD10.5 from 9-12 embryos 3 young dams and 6 9-12mo AAP dams. We analyzed the gene expression of pro-angiogenic *Vegfa, Vegfc,* and *Pgf* as well as the anti-angiogenic *sFlt1* – a soluble decoy receptor that binds VEGF family members, preventing their binding to signalling receptors.^56^ Decidual *sFlt1* expression was significantly decreased in the deciduae associated with developmentally delayed embryos, while *Vegfa, Vegfc,* and *Pgf* were unchanged between groups (Figure 4A). In the placenta there was an increase in *Vegfa* and *Pgf* expression in AAP compared to young, again driven by developmentally delayed embryos, with no change in *Vegfc* or *sFlt1* expression (Figure 4B). Thus, on both sides of the implantation site, angiogenic factor expression changes significantly with AAP, primarily driven by the status of the associated embryo.

**Figure 4:**
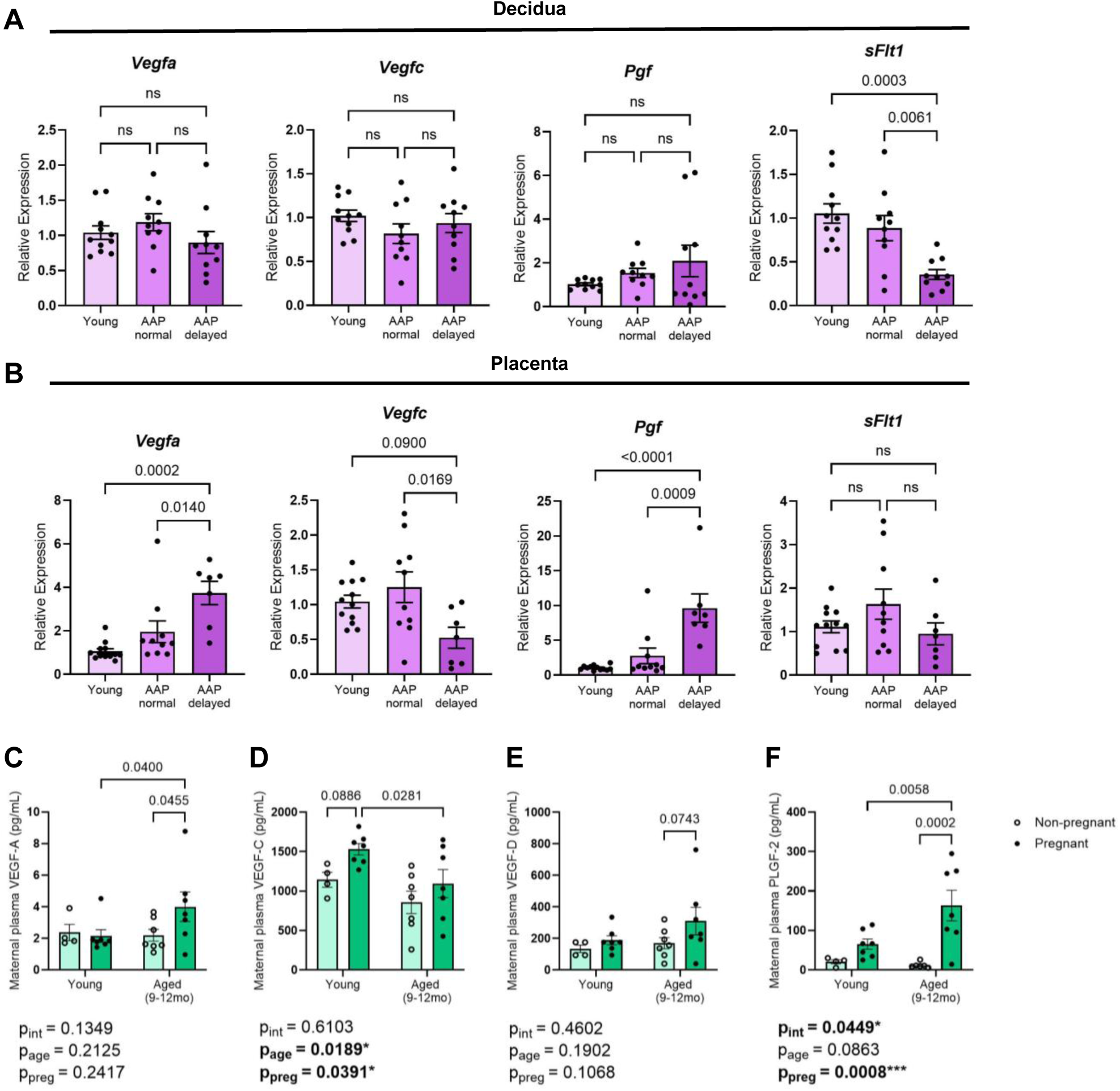
Circulatory and local VEGF angiogenic factor expression is altered in AAP. A-B) Summary data of the 2^−ΔΔCT^ in VEGF family gene expression in young vs AAP A) decidua and B) placenta from young embryos, developmentally normal AAP embryos, and developmentally delayed AAP embryos at GD10.5; C-F) Summary data of concentration of VEGF angiogenic factors (pg/mL) in maternal plasma in non-pregnant vs pregnant (GD10.5) young and aged (9-12mo) dams; A-B) mean ± SEM, one-way ANOVA with multiple comparisons; C-F) mean ± SEM, two-way ANOVA with multiple comparisons; 8-13 implantations from 4 young and 7 AAP dams were quantified.

Maternal plasma was collected from non-pregnant and GD10.5 young and aged (9-12mo) dams for multiplex analysis of circulating angiogenic factors. Of 16 probed pro-angiogenic factors and growth factors, 6 were significantly altered with pregnancy or age, of which 4 belonged to the VEGF family: VEGF-A (VEGF165 sub-isoform), VEGF-C, VEGF-D, and PLGF-2 (Figure 4C-F). While there was no change in circulating VEGF-A with pregnancy status in young mice, there was a significant pregnancy-specific increase in circulating maternal VEGF-A in the aged pregnant dams (Figure 4C). Notably, pregnancy significantly increased circulating VEGF-C in age-matched dams; however, age has a significantly opposing effect on circulating VEGF-C, causing VEGF-C levels to be significantly decreased in AAP vs young pregnant dams at GD10.5 (Figure 4D). Circulating VEGF-D was largely unchanged in young non-pregnant vs pregnant dams (Figure 4E). Circulating PLGF-2 increased within the pregnant aged group when compared to pregnant young and their age-matched non-pregnant littermates, but advancing age had an additive effect on this pregnancy-associated increase (Figure 4F). Further complexity can be observed when comparing young, aged (9-12mo), and aged (12+mo) mice (Supplementary Figure 5). Maternal circulating levels of VEGF factors are highly dynamic and show extensive alterations with age and pregnancy. Importantly, pregnancy-associated circulating VEGF-A and PLGF-2 levels are increased in AAP dams while pregnancy-associated circulating VEGF-C levels are diminished. In conjunction with decidual and placental expression of angiogenic factors, our data shows that expression of VEGF family members is imbalanced with AAP.

### VEGF-C is a pregnancy-associated angiogenic factor whose activation is altered in AAP decidua

One finding of particular interest is the loss of pregnancy-associated circulating VEGF-C in AAP. VEGF-C is a pro-angiogenic and pro-lymphangiogenic factor that binds to VEGFR2 and VEGFR3.^57^ VEGF-C binding affinity is improved through the sequential proteolytic cleavage of VEGF-C. Its final cleavage is mediated by ADAMTS3^57,58^ which is further enhanced by CCBE1.^58,59^ However, little is known concerning decidual VEGF-C localization, activation, and its role in decidual angiogenesis. When staining for anti-VEGF-C on young GD10.5 uterine sections, anti-VEGF-C signal is visible in both the placenta and decidua (Supplementary Figure 6). In the placenta anti-VEGF-C signal is visible in the stroma of the labyrinth zone and in trophoblast giant cells. In the decidua, anti-VEGF-C signal is concentrated in the vascular wall with considerable expression in the stroma. Colocalization of anti-VEGF-C signal with DBA^+^ uNK is consistent with findings by Pawlak *et al.* showing that uNK express VEGF-C in the murine decidua.^15^ Therefore, aside from the VEGF-C present in circulation, VEGF-C is present surrounding vessels in the decidua.

To better understand VEGF-C signalling in the uterine EC, we integrated uterine single cell RNA sequencing (scRNAseq) deposited datasets from Winkler *et al.* (2024)^60^ and Garcia Rivas *et al.* (2025).^61^ For these analyses, we used data from non-pregnant diestrus uteri, GD5.5 uteri, GD6.5 uteri, and GD10.5 decidua (Supplementary Table 1). Our integrated dataset had 82,904 cells including stromal cells, epithelial cells, immune cells (including NK cells, T cells, activated NK/T cells, dendritic cells, macrophages, mast cells), SMCs, and EC (Supplementary Figure 7). We further subclustered the EC, filtering for *Pecam1* positivity and *Ptprc* negativity, resulting in 764 cells (Figure 5A). As expected, expression of angiogenic receptors *Kdr* (VEGFR2), *Flt4* (VEGFR3), and *Tek* was largely exclusive to these EC along with other EC marker genes (Supplementary Figure 8, Supplementary Table 2). Within our EC subclusters, we had proliferative EC (*Mki67*+, *Pcna*+), activated EC (*Vcam1*+, *Icam1*+, *Sele*+, *Selp*+), *Pgr*+ EC, lymphatic EC (*Lyve1*+, *Prox1*+), arterial EC (*Efnb2*+, *Nr2f2*-, *Ephb4*-), and venous EC (*Nr2f2*+, *Ephb4*+) (Figure 5A, Supplementary Figure 9A-B). The proliferative EC subcluster was mostly comprised of EC from early gestation uterine tissue (Figure 5B), when Kim *et al.* (2013) documented significant proliferative vascular remodelling.^62^ Critically, we also have a cluster of remodelling spiral artery (SpA) EC that was contributed solely by cells from GD10.5 decidua (Figure 5B). Consistent with literature showing that spiral artery EC undergo intraendothelial transition,^15,63^ remodelling SpA EC express arterial markers (*Efnb2, Sox17*), venous markers (*Nr2f2, Ephb4*), and lymphatic marker *Lyve1* simultaneously (Supplementary Figure 9B-F).

**Figure 5:**
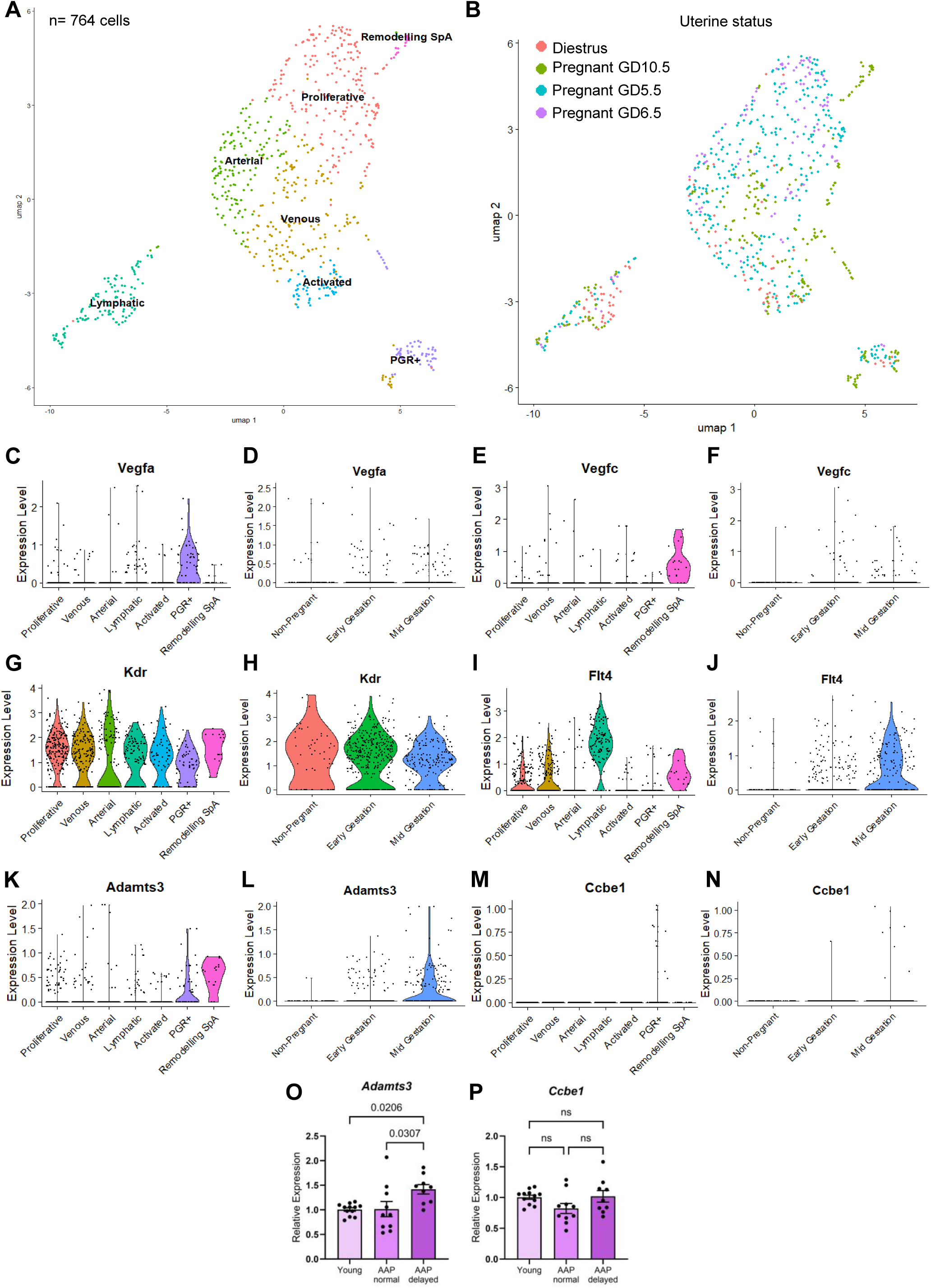
VEGF-C signalling is a potential endothelial autonomous signalling cascade that is altered in AAP. UMAP dimensionality reduction plot of endothelial cells subclustered from integrated young diestrus, aged diestrus, and young pregnant uterine scRNA-seq datasets grouped by A) subcluster identity and B) uterine status; Violin plots of C-D) *Vegfa*, E-F) *Vegfc,* G-H) *Kdr* (VEGFR2), I-J) *Flt4* (VEGFR3), K-L) *Adamts3*, and M-N) *Ccbe1* expression in endothelial cells based on C,E,G,I,K,M) endothelial cell subcluster identity and D,F,H,J,L,N) uterine status with lymphatic endothelial cells excluded; Summary data of the 2^−ΔΔCT^ in O) *Adamts3* and P) *Ccbe1* gene expression in decidua from young embryos, developmentally normal AAP embryos, and developmentally delayed AAP embryos at GD10.5; A-N) SpA = spiral arteries; O-P) mean ± SEM, student’s t-test; 9-12 implantations from 3 young and 7 AAP dams were assessed.

To examine VEGF signalling within these EC subclusters across gestation, particularly VEGF-C, we assessed the relative expression of *Vegfa, Vegfc, Kdr* (VEGFR2), *Flt4* (VEGFR3), *Adamts3*, and *Ccbe1* in these EC subclusters and across gestation (Figure 5C-N). For gestational analyses, lymphatic EC were excluded from the dataset. *Vegfa* expression was found to be most prominent within the PGR+ EC subcluster (Figure 5C) while *Vegfc* expression was most prominent within the remodelling SpA EC subcluster (Figure 5E), with select EC across other subclusters expressing these angiogenic factors. There were no notable changes in EC *Vegfa* expression across gestation (Figure 5D). While there was also no discernable change in the expression level of *Vegfc* across gestation (Figure 5F), stepwise increases in the percentage of *Vegfc+* EC with advancing gestation resulted in 15.6% of EC expressing *Vegfc* by mid-gestation (Supplementary Figure 10A).

As expected, *Kdr* (VEGFR2) was ubiquitously highly expressed across all EC subclusters (Figure 5G). Anti-VEGFR2 staining confirmed this ubiquitously expression across EC in various vessel types, with strongest expression at the lumenal surface (Supplementary Figure 11A). However, expression profiling captured a modest decrease in expression level of *Kdr* (VEGFR2) by mid-gestation (Figure 5H), despite no apparent change in the percentage of *Kdr*+ EC (Supplementary Figure 10B). *Flt4* (VEGFR3) was expressed highly in lymphatic, proliferative, venous, and remodelling SpA EC (Figure 5I). The elevated expression of *Flt4* (VEGFR3) observed in the proliferative EC suggests that VEGFR3 activity could be critical during EC proliferation in the early gestation period. Importantly, this also suggests that VEGF-C signalling affects multiple EC subpopulations across various vessel types. Anti-VEGFR3 staining shows that VEGFR3 was expressed on EC in vessels of various sizes (Supplementary Figure 11B). Contrary to *Kdr* (VEGFR2), *Flt4* (VEGFR3) positivity and relative expression within EC increased in a stepwise manner as pregnancy progressed (Figure 5J, Supplementary Figure 10C).

Lastly, when assessing the expression of genes required for proteolytic processing of VEGF-C, *Adamts3* was found to be highly expressed in select EC, most prominently within the PGR+ and remodelling SpA subclusters (Figure 5K). Similarly to *Flt4*, EC *Adamts3* expression increased in a stepwise manner as pregnancy progressed (Figure 5L). In the non-pregnant uterus only 1.6% of EC were *Adamts3+*, compared to 11.3% by early gestation, and 42.2% by mid-gestation (Supplementary Figure 10D). However, *Ccbe1* was not highly expressed in most EC (Figure 5M, N). The sum of our data suggests that EC can autonomously activate VEGF-A and VEGF-C signalling via angiocrine and paracrine interactions. Furthermore, scRNAseq analyses suggest that uterine VEGF-C signalling coordinates EC activity across gestation across multiple EC subtypes, increasing in prevalence by mid-gestation.

To quantitatively compare VEGF-C proteolytic activation between young and AAP deciduae, we used RT-PCR to assess *Adamts3* and *Ccbe1* expression in decidua from young implantations, AAP implantations with developmentally normal embryos, and AAP implantations with developmentally delayed embryos (Figure 5O-P). Decidual expression of *Adamts3* was upregulated in decidua from AAP implantations with developmentally delayed embryos compared to both young deciduae and AAP deciduae from developmentally normal embryos (Figure 5O). There was no change in the expression of *Ccbe1* (Figure 5P). This suggests that there is a transcriptional shift in the decidua towards increasing active VEGF-C abundance in AAP, potentially as a compensatory mechanism in developmentally delayed implantations.

### Decidual EC angiopoietin-2 expression is reduced in AAP

In our multiplex analyses of maternal plasma, we identified angiopoietin-2 (ANGPT2) as a circulating maternal non-VEGF angiogenic factor that is significantly increased with pregnancy; however, there was no compounding effect of age (Figure 6A). As an EC-derived autocrine, ANGPT2 binding to the TEK/TIE2 receptor plays key context-dependent roles in vascular destabilization, angiogenesis, vascular maturation, and responses to inflammation.^64–68^ Furthermore, it is known that the functional outcome of ANGPT2 is determined by the presence of VEGF.^67,69^ Aligning with previous literature,^70,71^ *Angpt2* and *Tek* expression in the uterus was restricted to endothelial cells (Supplementary Figure 8). When examining decidual and placental expression of *Angpt2,* contrary to our VEGF family findings, decidual *Angpt2* expression decreased regardless of embryo developmental status (Figure 6B). Compared to young placentae, AAP placentae from developmentally normal embryos had a trending increase in *Angpt2* expression that was lost in AAP developmentally delayed embryos (Figure 6B). Thus, in addition to alterations in VEGF signalling, our data also suggests alterations in ANGPT2 signalling with AAP. However, unlike our VEGF-C findings this impairment is not present in systemic maternal circulation but instead localized to the implantation bed.

**Figure 6:**
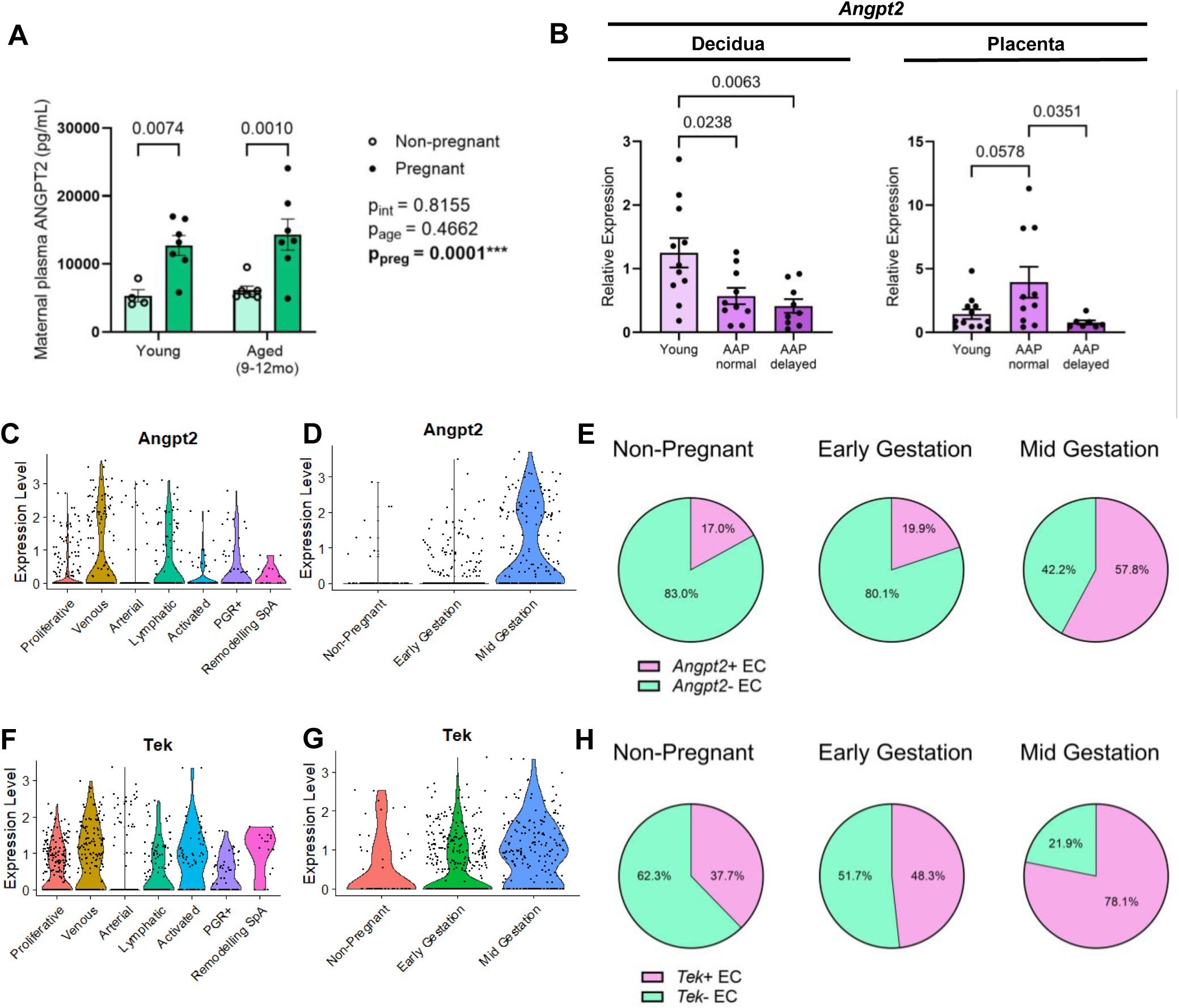
Angiopoietin 2 is an endothelial autocrine that is impaired in AAP decidua. A) Summary data of concentration of ANGPT2 (pg/mL) in maternal plasma in non-pregnant vs pregnant (GD10.5) young and aged (9-12mo) dams; B) Summary data of the 2^−ΔΔCT^ in *Angpt2* expression in young vs AAP decidua and placenta from young embryos, developmentally normal AAP embryos, and developmentally delayed AAP embryos at GD10.5; Violin plots of C-D) *Angpt2* and F-G) *Tek* expression in endothelial cells based on C,F) endothelial cell subcluster identity and D, G) uterine status; E, H) Summary data of the percentage of E) *Angpt2+* and H) *Tek+* endothelial cells over all endothelial cells; A) mean ± SEM, two-way ANOVA with multiple comparisons; B) mean ± SEM, one-way ANOVA with multiple comparisons; D,E,G,H) Lymphatic endothelial cells were excluded from these analyses.

To better understand decidual ANGPT2 signalling during pregnancy, we assessed the expression of *Angpt2* and its receptor *Tek* across EC subclusters and across gestation using our integrated dataset (Figure 6C-H). *Angpt2* was widely expressed across multiple EC subclusters, particularly within venous, lymphatic, and PGR+ EC (Figure 6C). With advancing gestation, vascular *Angpt2* expression (Figure 6D) and percentage of *Angpt2+* EC increased (Figure 6E); however, unlike VEGF signalling mediators this did not occur in a stepwise fashion. Relative EC expression of *Angpt2* increased dramatically between early and mid-gestation (Figure 6D). Additionally, between non-pregnant and early gestation uterine EC, there was a modest increase of 17.0% to 19.9% in *Angpt2+* EC, but by mid-gestation, *Angpt2+* EC increased to 57.8% of all EC (Figure 6E). Expression of the ANGPT2 receptor *Tek* was also widely observed across multiple EC subclusters, except for arterial EC (Figure 6F). Relative *Tek* expression and percentage of *Tek+* EC increased across gestation in a stepwise manner (Figure 6G, H). In summary, our data shows that autocrine ANGPT2 signalling is enriched in decidual EC at mid-gestation. Furthermore, AAP decidua at GD10.5 regardless of embryonic outcome have decreased *Angpt2* expression compared to young, revealing that there is localized decidual endothelial impairment in AAP.

## Discussion

With increasing maternal age there is a well-documented increase in risk of adverse pregnancy outcomes such as preeclampsia, intrauterine growth restriction, gestational hypertension, and stillbirth.^21–25^ Transformation of the uterine lining into the decidua occurs alongside angiogenic decidual vascular expansion and remodelling to mediate maternal adaptations to pregnancy. We found that in conjunction with poor embryonic outcomes and placental malformations, AAP mice also exhibit significantly impaired decidual angiogenesis, impaired spiral artery remodelling, and imbalances in angiogenic signalling both systemically and within the implantation site. Impaired spiral artery remodelling has been broadly accepted as a foundational driver of preeclampsia and associated fetal growth restriction.^17,18,72^ Our findings that spiral artery remodelling was likewise impaired in AAP suggests that this defect is more prevalent with age and may contribute to heightened pregnancy complication risk of particularly preeclampsia, aligning with epidemiological studies.^21–25^ Given the severely elevated lifetime maternal cardiovascular disease risk posed by a pregnancy complicated by preeclampsia,^73^ it is possible that age-associated defects in uteroplacental perfusion and secondary systemic effects may contribute to lifelong cardiovascular risk in AAP parents.

These profound impairments in decidual angiogenic vascular remodelling with AAP have not been shown before. Previously, the non-pregnant uterus has been reported as a tissue that does not exhibit vascular attrition with age.^34^ It was previously hypothesized that this characteristic was due to its highly regenerative nature; however, our data suggests that age-associated vascular attrition does occur, but it may be restricted to or exacerbated by pregnancy. Furthermore, the sum of our analyses suggests that in addition to EC extrinsic aging, EC intrinsic deficiencies and refractory vasculature are key features of AAP. Consistent with this, AAP deciduae had reduced EC abundance, vascular density, and a shift towards small calibre vessels across all vessel types. Critically, at mid-gestation endothelial proliferation is unchanged between young and AAP decidua, suggesting that any AAP-associated disruptions in proliferative decidual angiogenesis occur earlier in gestation. However, this does not discount the possibility that non-proliferative mechanisms of angiogenesis and vessel widening, such as EC migration, are disrupted at GD10.5. Hence, the double hit of age and necessity for expansion of the vascular network reveal an aging phenotype consistent with other tissues. Thus, the vascular effects of aging in the uterus may not be fully unmasked until the introduction of pregnancy. We posit that the aging decidua is unable to appropriately respond, resulting in poor vascular remodelling.

Despite different vascular origins, the anatomical proximity and shared interface of the decidua and placenta enable early maternal-fetal communication. Murine models have shown that poor expansion of decidual vasculature is associated with poor fetal capillary expansion.^74,75^ Interestingly in this study, although both placental and decidual vascularity is decreased in AAP dams, vascularity of one is independent of the other at mid-gestation. However, the relationship between decidual and placental vascularity at other gestational timepoints is not known. Utero-placental blood flow in mice commences at GD10.5 and increases progressively with advancing gestation.^46–48^ Furthermore, both decidual and placental vascular remodelling continue beyond GD10.5 and through GD15.5.^11,76,77^ Given the importance of shear stress in shaping endothelial function and angiogenic signalling,^78–81^ there may be a growing interdependence between the decidual and placental vasculature as utero-placental blood flow increases.

In this study, we identified AAP-associated imbalances in the expression of VEGF and angiopoietin signalling mediators, including decreases in systemic VEGF-C, increases in both systemic and placental VEGF-A and PGF, and decreases in decidual sFlt1 and ANGPT2. Excesses of pro-angiogenic factors, like VEGF-A and PGF, are not necessarily conducive to pregnancy success. This is further complicated by decidual-placental communication. Although increasing systemic VEGF-A has been shown to prevent vascular attrition and prolong the lifespan of aging mice,^39^ endometrial VEGF-A supplementation has adverse effects on pregnancy outcome including embryo loss, placental vascular malformations, and upregulation in placental sFlt1.^54^ Placental VEGF-A deletion has been shown to increase circulating maternal VEGF-A and negatively affect maternal cardiac function.^55^ Hence, a balance of angiogenic factors between the decidua and placenta is necessary for pregnancy progression and maintenance of maternal cardiovascular function. Additionally, given the pleiotropic and interactive nature of VEGF and angiopoietin signalling,^67,69^ these imbalances are likely synergistic, producing a sum effect irreproducible by one factor alone. Ultimately, whether these imbalances in angiogenic signalling contribute to the impaired decidual and placental vascularization or reflect an acute compensatory response to poor vascularization remains unknown. Future studies will be needed to identify the stage at which decidual angiogenic signalling is impaired and how progression of decidual angiogenesis in AAP dams may differ from young.

Importantly, we identify VEGF-C signalling as an area of interest in AAP. In pregnancy, VEGF-C knockdown in hematopoietic and endothelial lineages (via the Vav1-Cre transgenic line) impair spiral artery remodelling;^15^ however, little is known concerning VEGF-C’s role in preceding stages of decidual angiogenesis. Blocking VEGFR2 or VEGFR3 signalling during pregnancy leads to impaired vascular remodelling; but only blockage of VEGFR2 precludes pregnancy progression.^7,8,15^ Thus, it is possible that any impairments in VEGF-C/VEGFR3 activity during pregnancy is partially compensated for by the activity of VEGFR2. However, VEGFR3 may play a key role in further tuning decidual angiogenesis. It has been shown in zebrafish and mice that endothelial autonomous signalling via VEGF-C is essential in both angiogenesis and lymphangiogenesis.^82,83^ Our integrated scRNAseq analyses likewise suggests that EC-autonomous VEGF-C/VEGFR3 interactions may play key roles in the uterine endothelium during pregnancy. Further investigation into VEGF-C in AAP, its systemic sources in pregnancy, and its potential as a key angiocrine during decidualization is necessary.

We have also identified ANGPT2 as a decidual angiocrine that is altered with AAP. ANGPT2 abundance increases with pregnancy^70,71,84^ and its expression is restricted to EC in both the human and murine pregnant uterus.^70,84^ Our scRNAseq analyses shows that EC autonomous *Angpt2* signalling is elevated specifically at mid-gestation, suggesting that ANGPT2 signalling is particularly important at this point in gestation.

Furthermore, the decreased expression in decidual *Angpt2* without alteration in maternal circulating ANGPT2 and regardless of embryonic outcome implies that there is an isolated impairment localized within decidual EC in AAP. However, the effect of placental ANGPT2 on the maternal decidua is unknown. Placental upregulation of *Angpt2* in developmentally normal embryos may act as a compensatory mechanism, but whether this compensates for deficient placental vascularization or deficient decidual vascularization is unknown.

In summary, we show that decidual angiogenic vascular remodelling is impaired in a murine model of AAP. AAP dams have significantly altered systemic, decidual, and placental expression of angiogenic factors. It has been shown in other tissues that vascular attrition and VEGF signalling insufficiency are features of aging. Yet a paucity of data on the aging uterine vasculature and its adaptation to pregnancy hinders our understanding of AAP or how we can alleviate age-associated risks during pregnancy.

## Materials and Methods

### Study approval

The study was conducting in accordance with the procedures outlined in the animal ethics (AUP00004423) approved by the University of Alberta animal ethics committee and in accordance with the Canadian Council on Animal Care guidelines.

### Animal model

C57BL/6J mice (Jackson Laboratories) were bred at the University of Alberta conventional mouse facilities. Female virgin C57BL/6J mice were aged to 11-16wk or 40-70wk. Aged cohorts, starting at 20wk, were pair-fed. Young and AAP mice were mated with young (11-16wk) C57BL/6J males and GD0.5 was determined by presence of a copulation plug the following morning.

Dams at GD10.5 were euthanized using cardiac puncture under anesthesia with isoflurane, followed by cervical dislocation. Maternal plasma was stored with EDTA and flash frozen. Uterine horns were either placed in ice-cold PBS for further dissection of the embryos, placentae, and deciduae or fixed in 4% paraformaldehyde overnight for downstream histology and immunostaining. Embryos were dissected and imaged using a Zeiss Stemi 508 stereomicroscopy. Isolated placentae and deciduae were flash frozen for downstream RNA isolation. See Supplemental Methods for details.

### Histology, immunohistochemistry, lectin, and immunofluorescence staining

Mid-sagittal serial 10μm cryosections were sectioned from uterine horns then stained with H&E (Sigma-Aldrich), Masson’s Trichrome (Polysciences), or immunostained with primary and secondary antibodies detailed in Supplementary Table 3. For immunohistochemistry, placental trophoblasts were detected using anti-pan-cytokeratin and uNK cells with DBA-lectin then developed using a DAB peroxidase substrate kit (Vector Labs). For immunofluorescence smooth muscle cells were detected with anti-α-SMA, EC were detected with anti-CD31 and anti-Erg1/2/3, and Hoechst 33342 was used as a nuclear dye. See Supplemental Methods for details and product info.

### Imaging and image analysis

Sections were imaged using a Zeiss Axioscan.Z1 slide scanner at 10X or with a Zeiss LSM 700 confocal microscope at 10X and 20X. Images were analyzed using ZEN blue 3.6 (Zeiss) and Volocity 7.0.0 (Quorum Technologies) imaging softwares. Vessels were traced using Vesselucida 360 2022.1.1 (MBF Bioscience) and quantified using Vesselucida Explorer 2022.1.1 (MBF Bioscience). See Supplemental Methods for details and product info.

### RNA isolation and RT-PCR

Total RNA from deciduae or placentae was isolated using TRIzol-chloroform (Thermo Fisher Scientific) extraction and purified using a PureLink RNA Mini Kit (Ambion). 500ng of RNA was converted into cDNA using an iScript cDNA Synthesis Kit (Biorad). RT-PCR was performed using an Applied Biosystems SYBR Green PCR Master Mix (Thermo Fisher Scientific) on a QuantStudio Real-Time PCR System (Thermo Fisher Scientific) as previously described.^85^ For primer sequences see Supplementary Table 4. See Supplemental Methods for details and product info.

### Integration and analysis of scRNAseq

Filtered datasets were used from Winkler *et al.* (2024) EMBL-EBI ArrayExpress accessions E-MTAB-11491, E-MTAB-12889 and from Garcia Rivas *et al.* (2025) NCBI GEO accession GSE299135. For sample information of the deposited datasets utilized, see Supplementary Table 1. All scRNAseq data was analyzed and visualized in Seurat 5.4.0 in R 4.5.2 and R Studio 2026.01.1+403. Integration and batch correction for samples was performed using CCA integration. Within our 82,904 cells, we identified subclusters using various marker genes. For marker genes used to identify subclusters within the integrated Uniform Manifold Approximation Projection (UMAP), see Supplementary Table 2. We subset our EC by filtering *Pecam1* > 0.1 and *Ptprc* = 0. These EC subsets were further integrated and batch corrected using CCA integration.

### Statistics

Data was analyzed using unpaired parametric and non-parametric student t-test, one-way ANOVA, or two-way ANOVA with multiple comparisons. All tests performed were two-sided and tested for normality. Categorical data was analyzed using a chi-squared test or Fisher’s exact test. P<0.05 was determined statistically significant. Analysis was performed using Graphpad Prism 10.4.2.

To assess the ability of placental and decidual vascularity to predict one another, linear mixed effects models were constructed in R 4.6.0 using the lmerTest package^86^ A reduced additive model was selected based on a lack of significant interaction between maternal age and vascular fraction measurement. Additive models were analyzed using ANOVA with Type II sum of squares and Satterthwaite calculated denominator degrees of freedom. Model assumptions of residual normality, homoscedasticity, conditional independence and the absence of multicolinearity was performed using a combination of base ‘plot’ functions and acf, vif, and qqplot functions from the car package 3.1.5.^87^

## Acknowledgements

We would like to acknowledge the salary support for W.K.D. from the Canadian Institutes of Health Research and One Child Every Child, and the expert assistance of the University of Alberta North Campus Animal Services for animal husbandry.

## Author contributions

M.R., W.K.D., and J.G.R. designed the studies. W.K.D., N.N., A.S., E.C., C.B., and J.G.R. conducted experiments, acquired data, and analyzed data used in this study.

Bioinformatic analyses were performed by W.K.D. W.K.D. prepared figures and wrote the manuscript. All authors revised drafts and reviewed the manuscript.

## Data Availability

All reagents and materials used in this study are available in Supplementary Methods and Materials and in Supplementary Tables 3-4. Publicly deposited scRNA-seq data used in this study are available in Supplementary Table 1, in ArrayExpress at accession numbers E-MTAB-12889, E-MTAB-11491, and in Gene Expression Omnibus (GEO) at accession number GSE299135. Source data behind the figures in this paper are available in the Supplemental data file. Any remaining information can be obtained from the corresponding author upon reasonable request.

