## Supplementary Figures for "Advanced Maternal Age Impairs Decidual Angiogenesis and Spiral Artery Remodelling"

**Supplemental Figure 1**

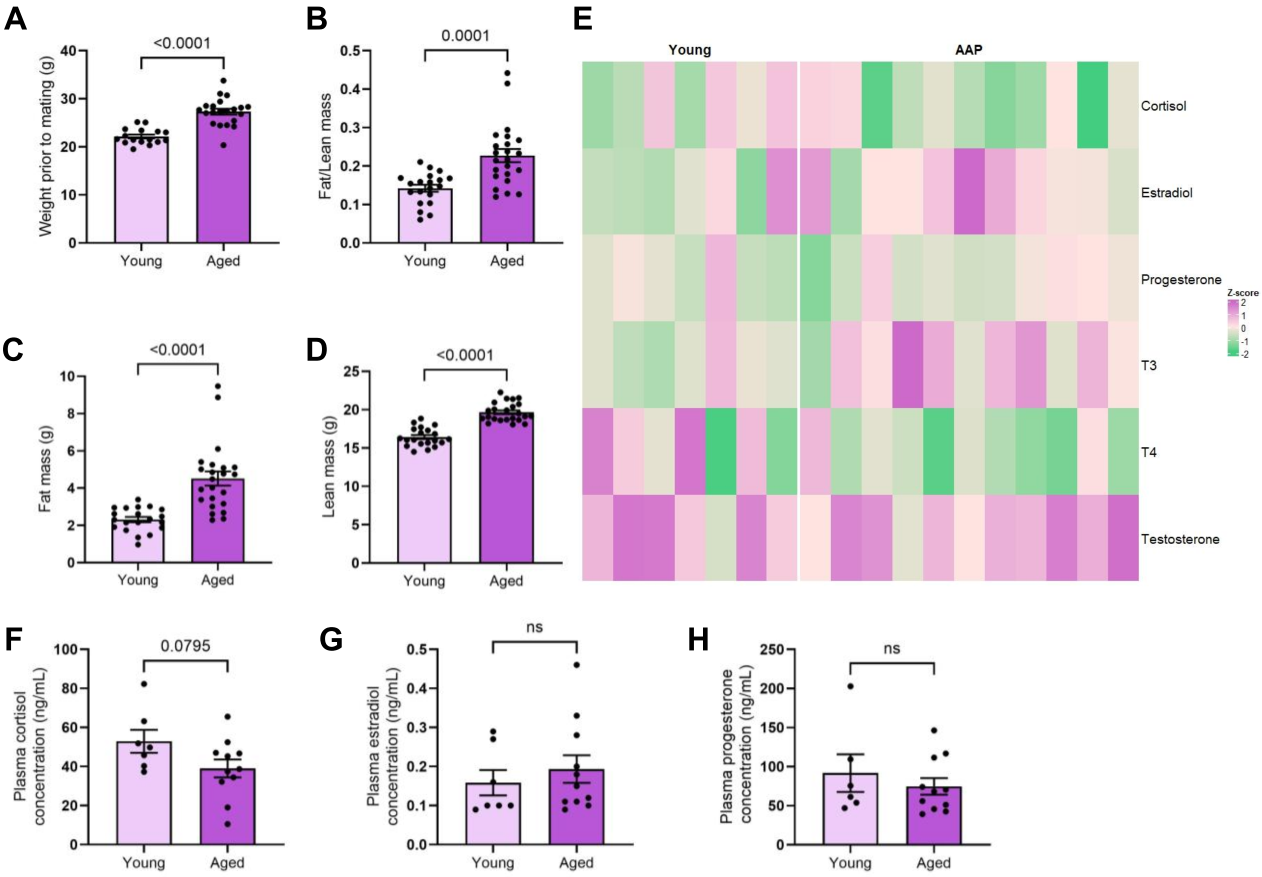


**Supplemental Figure 1: Maternal characteristics of aged female mice compared to young.** Summary data of A) maternal weight prior to mating (g), B) fat:lean mass, C) fat mass (g), and D) lean mass (g) prior to mating; E) Heatmap of circulating steroid hormones at GD10.5 by z-score; Summary data of concentration of circulating F) cortisol (ng/mL), G) estradiol (ng/mL), H) progesterone (ng/mL) in maternal plasma at GD10.5; A-D), F-H) mean ± SEM, student’s t-test.

**Supplemental Figure 2**

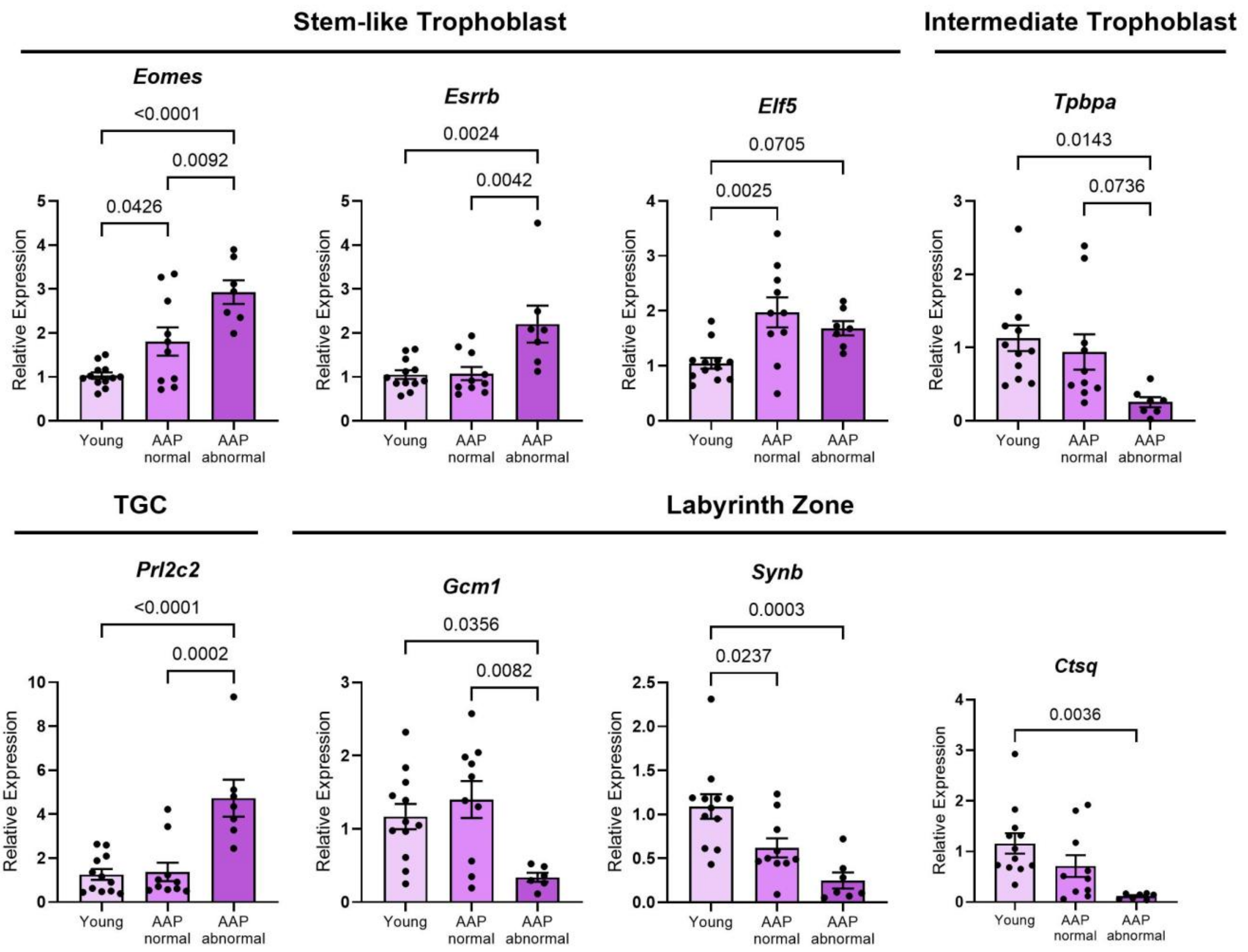


**Supplemental Figure 2: Significant alterations in placental composition at GD10.5 with AAP.** Summary data of the $2^{-\Delta\Delta CT}$ in placental marker gene expression of placentae from young embryos, developmentally normal AAP embryos, and developmentally delayed AAP embryos at GD10.5; mean ± SEM, one-way ANOVA with multiple comparisons.

**Supplemental Figure 3**

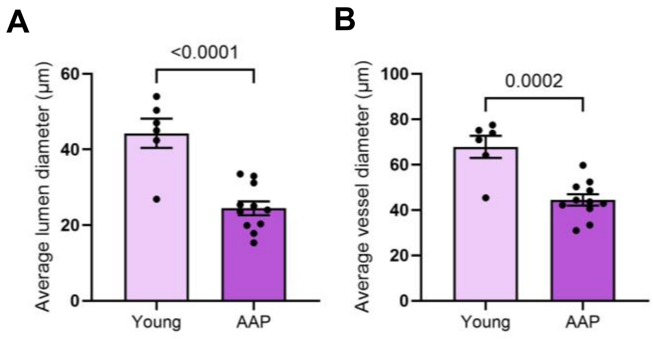


**Supplemental Figure 3: Spiral arterial dilation at GD10.5 is impaired in AAP decidua.**

Summary data of the average A) lumen diameter (µm) and B) vessel diameter (µm) of spiral arteries in young vs AAP decidua; mean ± SEM, unpaired parametric student's t-test; 6-11 implantations from 3 young and 4 AAP dams were quantified.

### Supplemental Figure 4

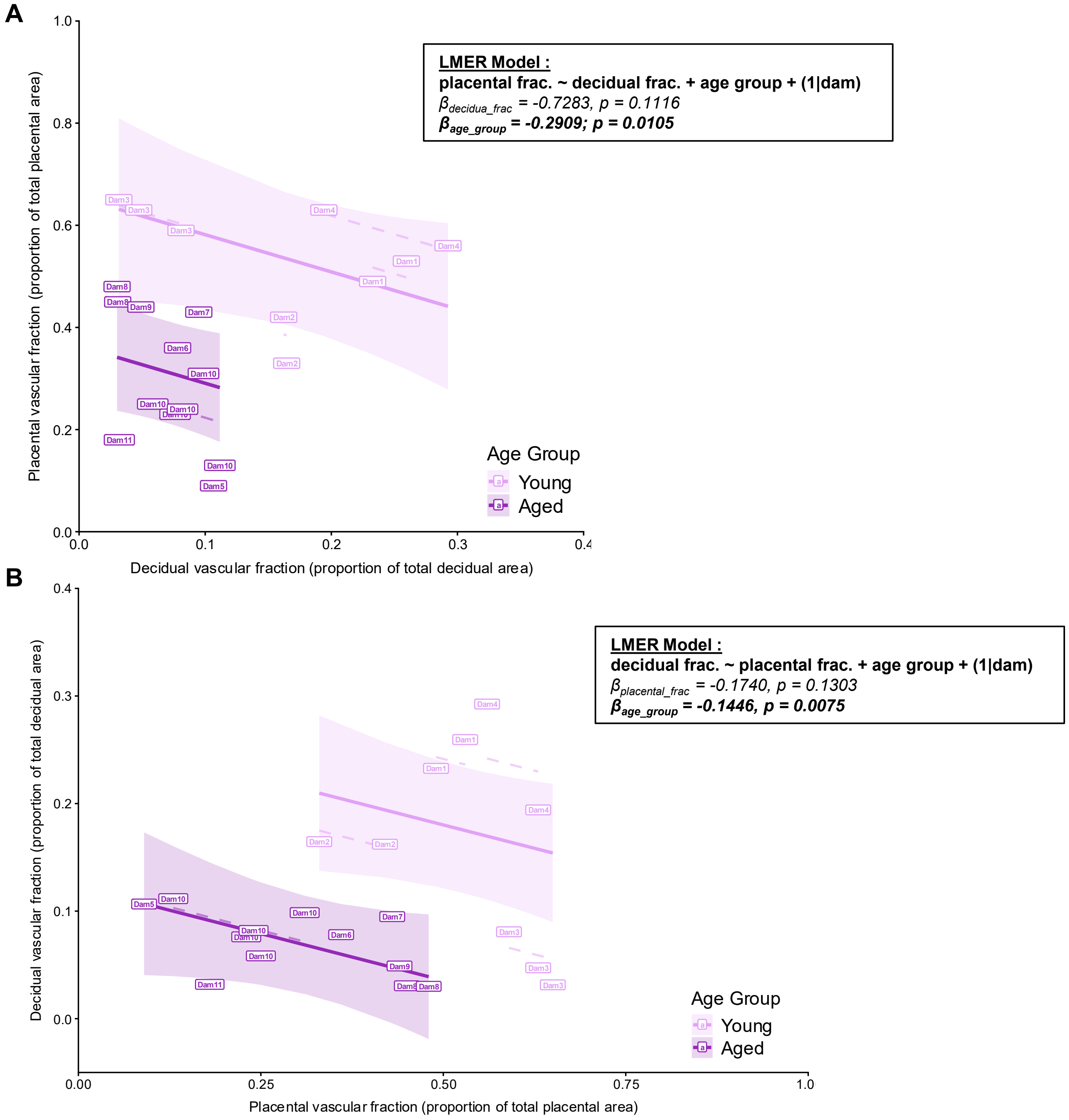


**Supplemental Figure 4: Vascularity of the decidua is not a statistically significant predictor of placental vascularity and vice versa at GD10.5.** Linear mixed-effects regression analyses of A) placental vascularized fraction as a function of decidual vascular density and maternal age and B) decidual vascular density as a function of placental vascularized fraction and maternal age, nested by dam.

**Supplemental Figure 5**

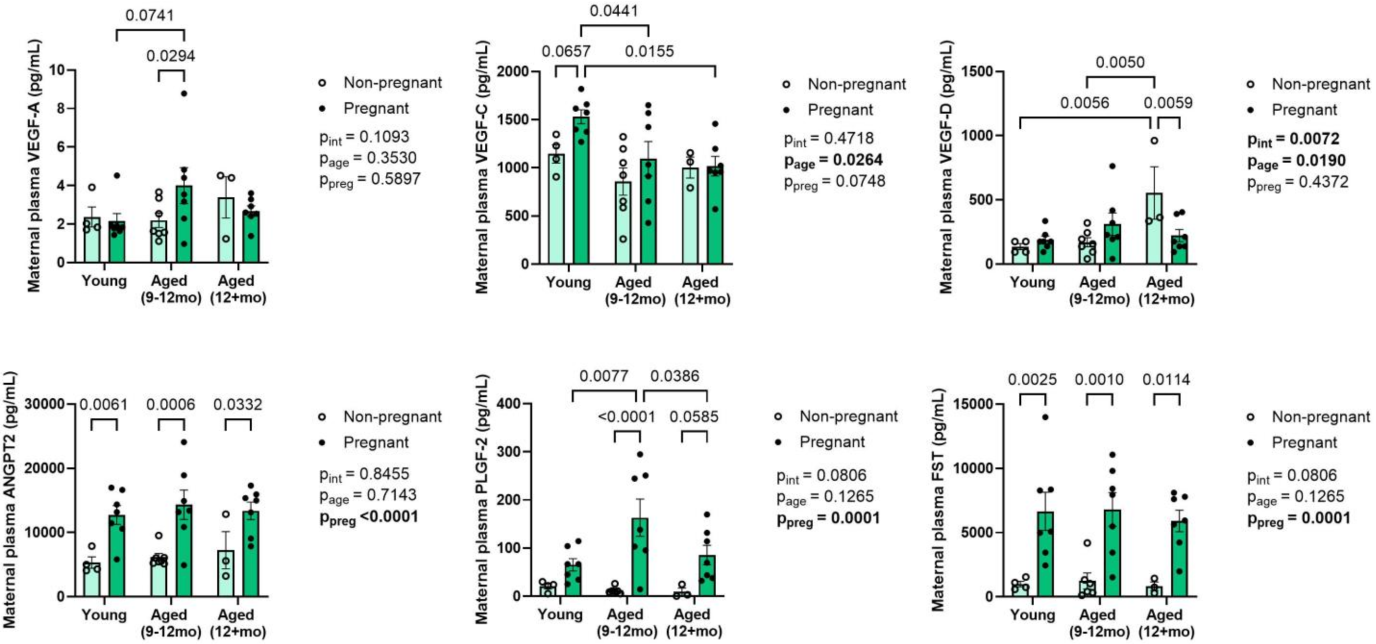


**Supplemental Figure 5: Maternal circulating angiogenic factor expression is altered in AAP.** Summary data of concentration of VEGF-A (pg/mL), VEGF-C (pg/mL), VEGF-D (pg/mL), ANGPT2 (pg/mL), PLGF-2 (pg/mL), and FST (pg/mL) in maternal plasma in non-pregnant vs pregnant (GD10.5) young, aged (9-12mo), and aged (12+mo) dams; mean ± SEM, two-way ANOVA with multiple comparisons; 7-12 implantations from 3 young and 7 AAP dams were quantified.

**Supplemental Figure 6**

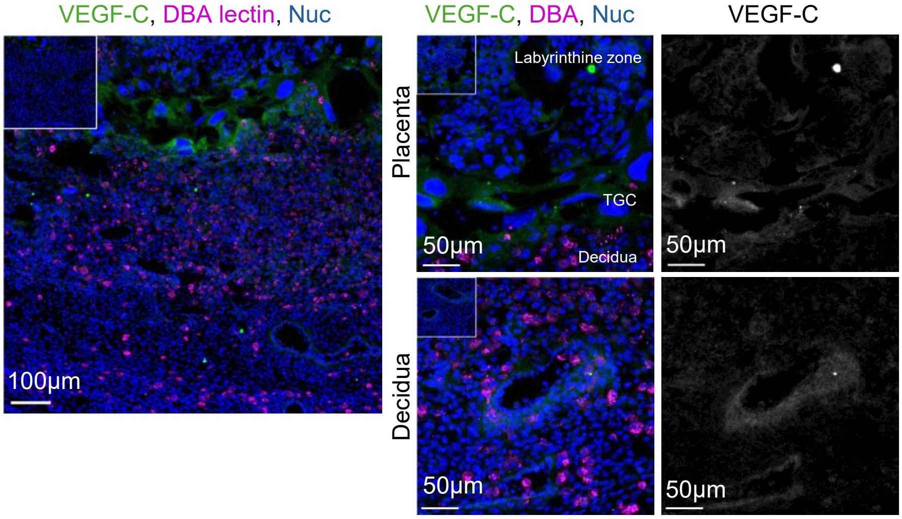


**Supplemental Figure 6: VEGF-C is localized within the vessel wall of the decidua and within the placenta.** Representative confocal microscopy of a murine implantation site stained for anti-VEGF-C, DBA lectin (uNK cells) and Hoechst in the placenta and the decidua; insets = isotype negative controls of respective areas stained with Hoechst.

### Supplemental Figure 7

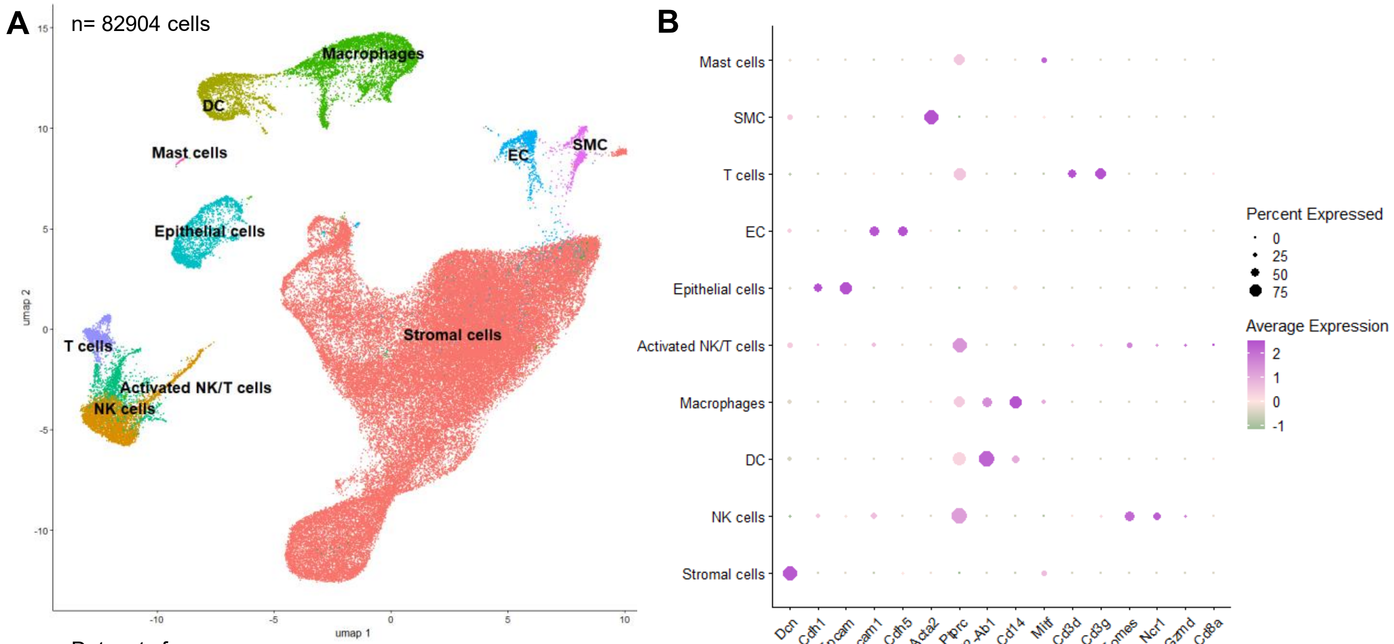


Datasets from:
Winkler *et al.* Cell. 2024.
Garcia Rivas *et al.* Biol. Reprod. 2025.

**Supplemental Figure 7: Integrated uterine scRNAseq dataset consisting of non-pregnant diestrus, GD5.5, GD6.5, and GD10.5 C57BL/6J uteri.** A) UMAP dimensionality reduction plot of 82904 cells clustered from integrated young diestrus, aged diestrus, and young pregnant uterine scRNA-seq datasets from Winkler *et al.* (2024) and Garcia Rivas *et al.* (2025); B) Dot plot showcasing the average expression and percentage of cells expressing key marker genes across all identified uterine subclustered cell types.

**Supplemental Figure 8**

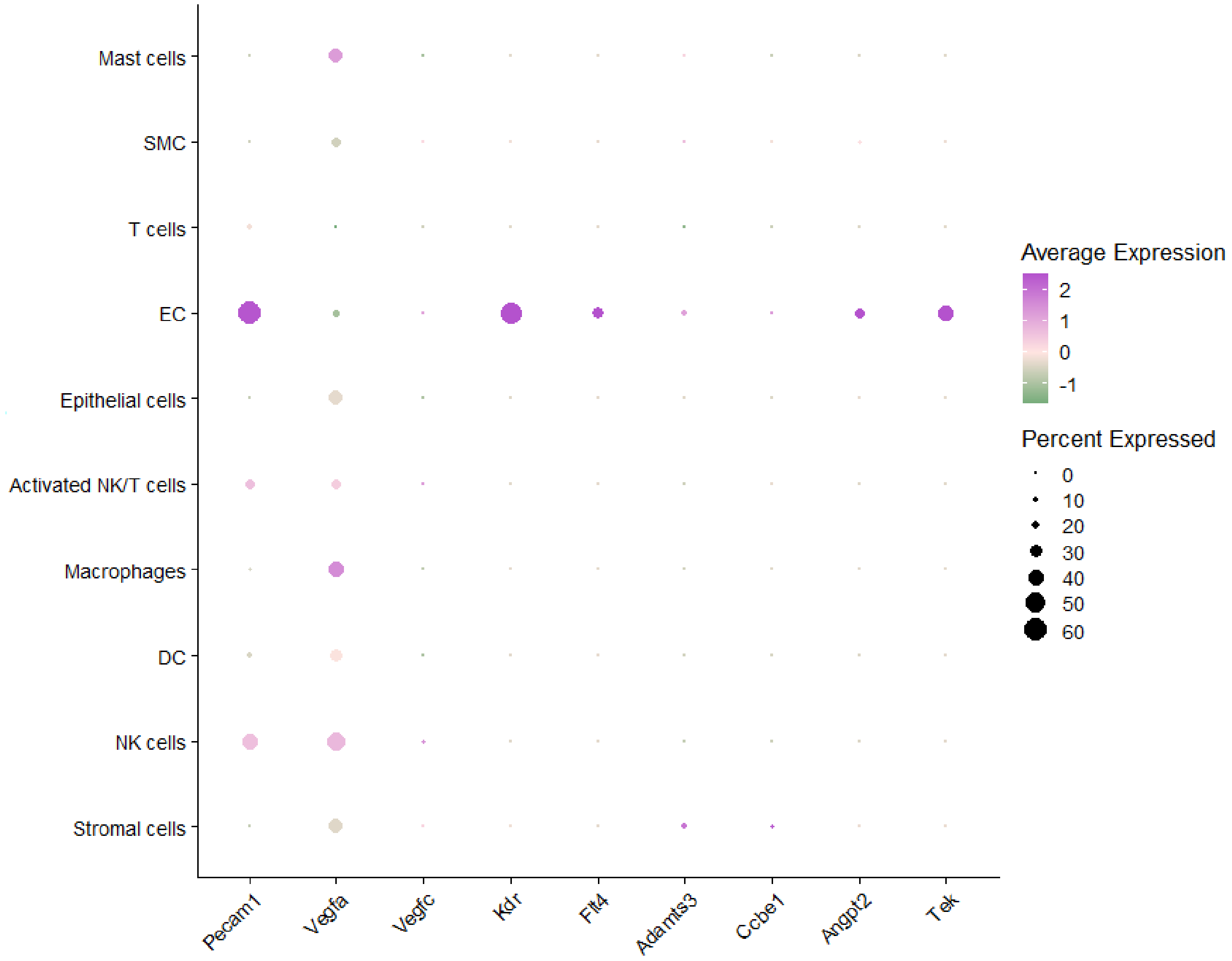


**Supplemental Figure 8: Expression of angiogenic factors and receptors across uterine cell types.** Dot plot showcasing the average expression and percentage of cells expressing *Pecam1, Vegfa, Vegfc, Kdr, Flt4, Adamts3, Ccbe1, Angpt2,* and *Tek* across all identified uterine subclustered cell types.

**Supplemental Figure 9**

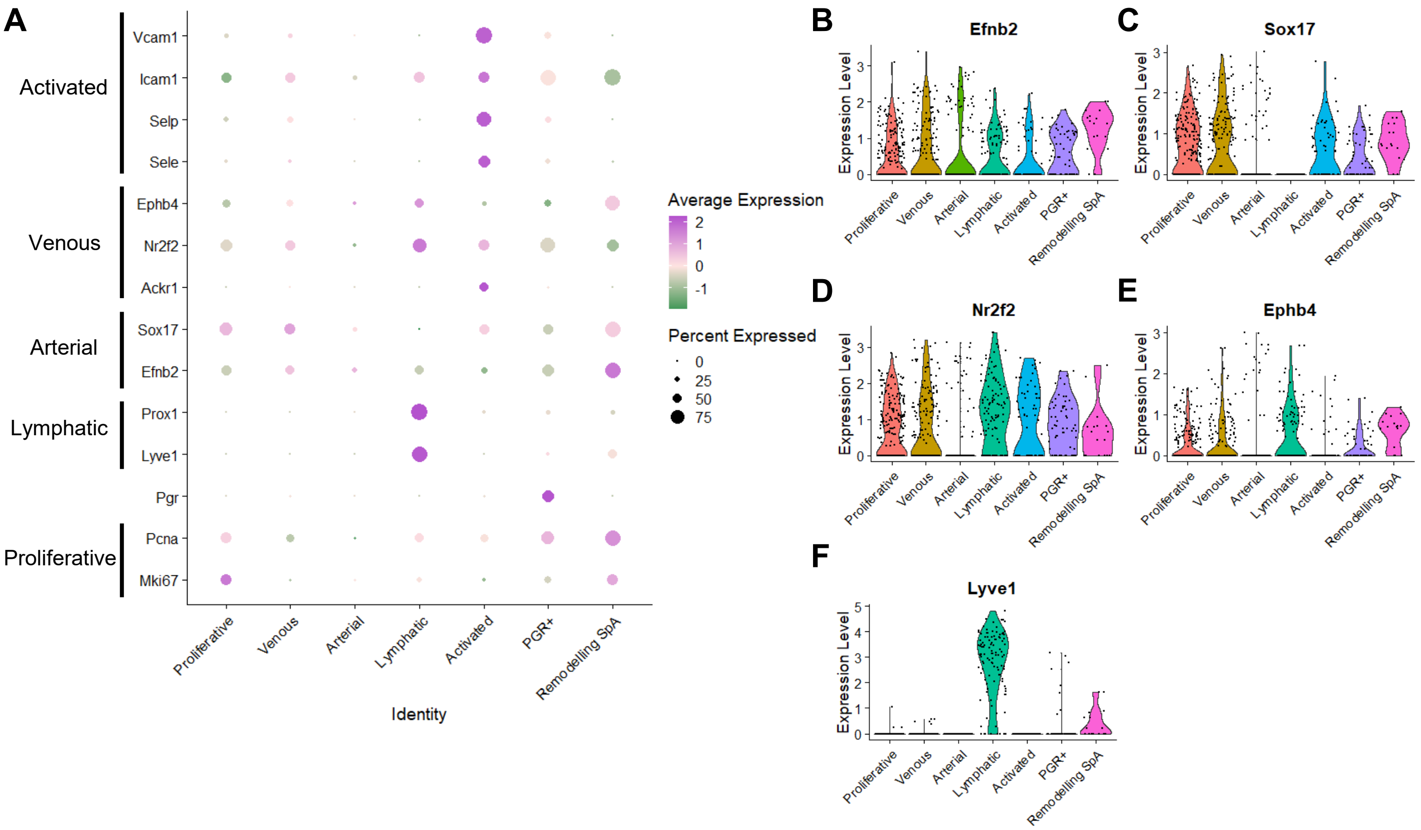


**Supplemental Figure 9: Identification of EC subclusters.** A) Dot plot showcasing the average expression and percentage of cells expressing marker genes associated with EC subcluster identity; B-F) Violin plots showcasing the expression of arterial markers B) *Efnb2* and C) *Sox17,* venous markers D) *Nr2f2* and E) *Ephb4,* and lymphatic marker F) *Lyve1* across EC subcluster identities.

**Supplemental Figure 10**

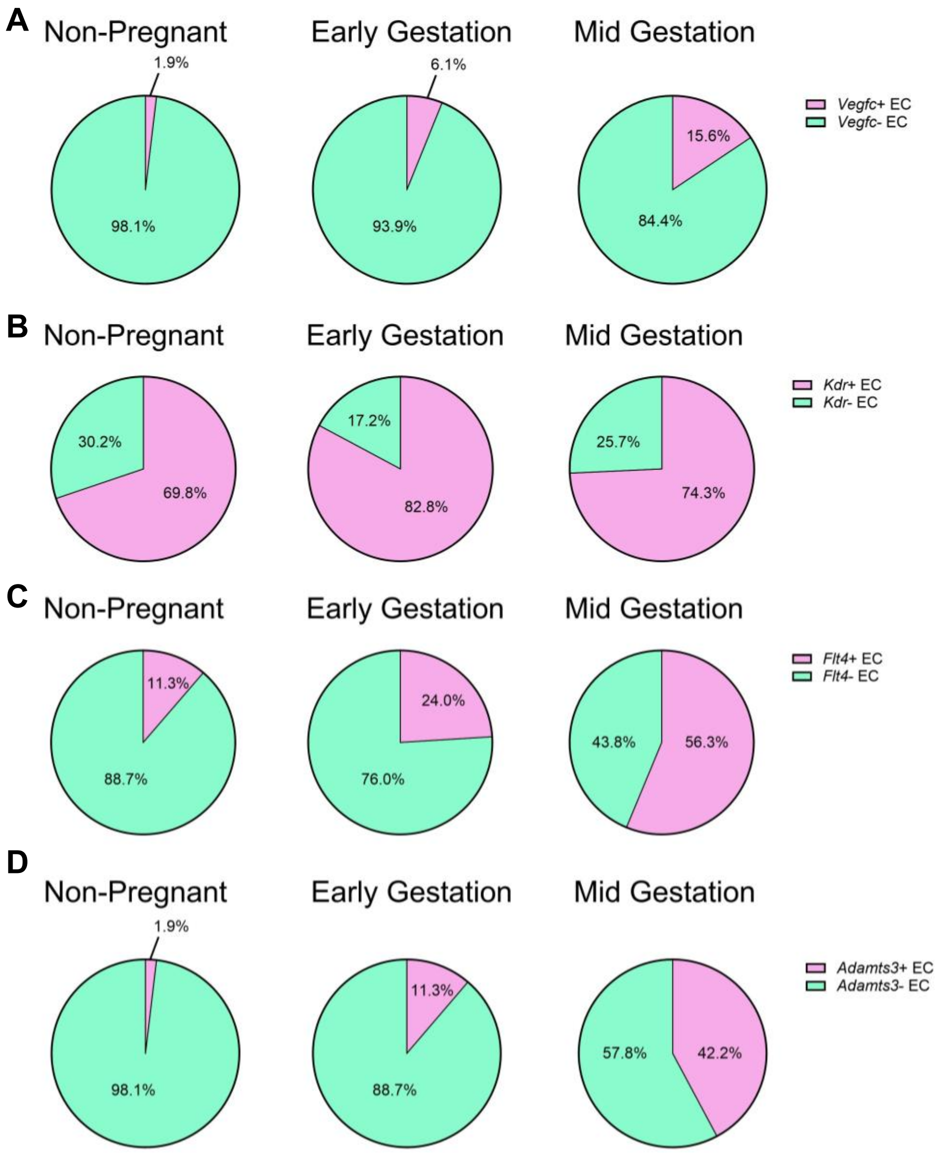


**Supplemental Figure 10: Percentage of EC expressing VEGF-C signalling mediators increases in a stepwise manner from non-pregnant towards mid-gestation.** Summary pie charts showing the percentage of A) *Vegfc*+ and *Vegfc*- EC, B) *Kdr*+ and *Kdr-* (VEGFR2) EC, C) *Flt4*+ and *Flt4-* (VEGFR3) EC, and D) *Adamts3*+ and *Adamts3*- EC.

### Supplemental Figure 11

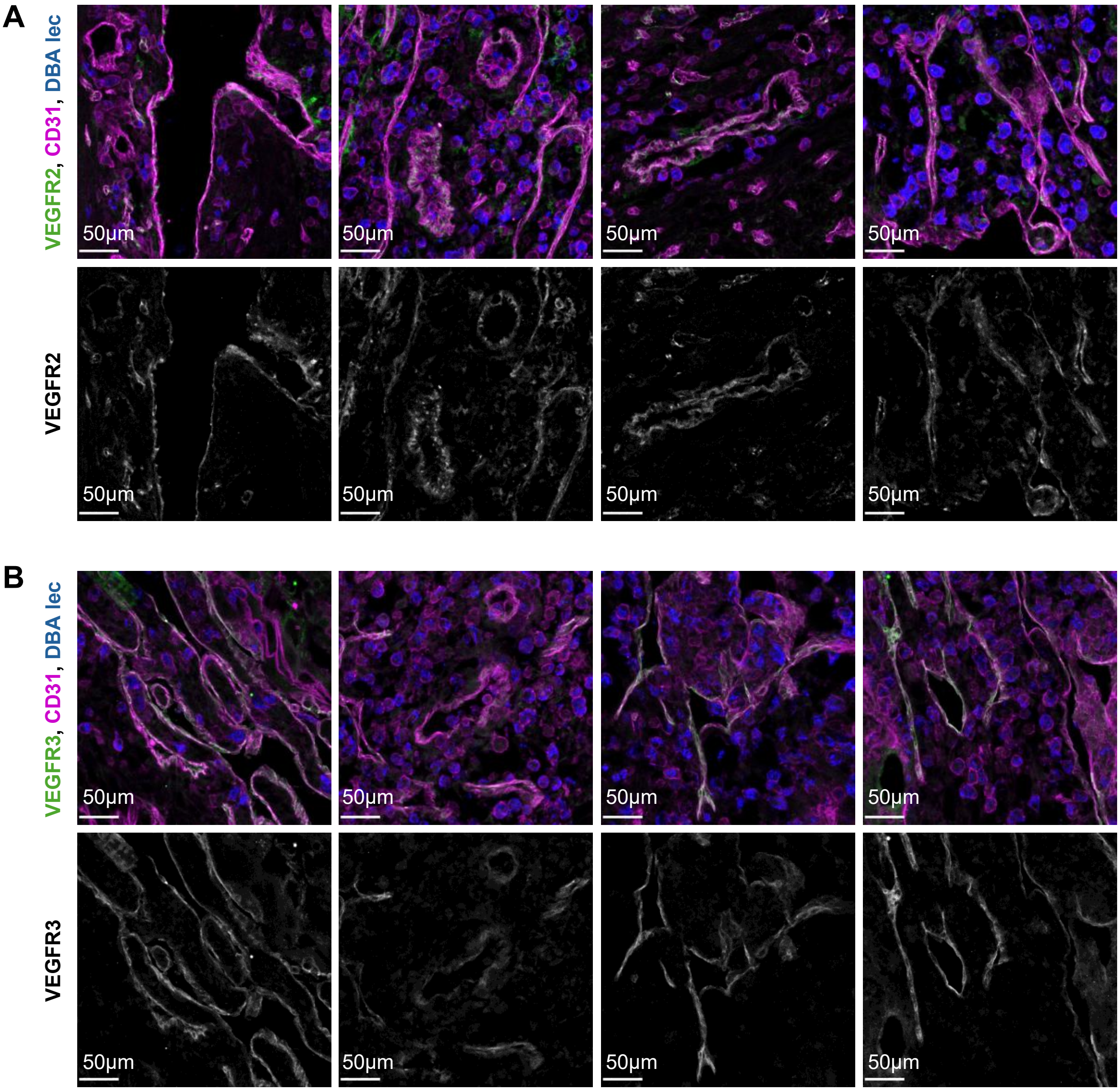


**Supplemental Figure 11: VEGFR2 and VEGFR3 expression across decidual EC.**

Representative confocal microscopy images of decidual vessels stained for anti-CD31, DBA lectin (uNK cells), and A) anti-VEGFR2 or B) anti-VEGFR3.
